# A genomic atlas of the human gut virome elucidates genetic factors shaping host interactions

**DOI:** 10.1101/2025.11.01.686033

**Authors:** Antonio Pedro Camargo, Fotis A. Baltoumas, Eric Olo Ndela, Mateus B. Fiamenghi, Bryan D. Merrill, Matthew M. Carter, Yishay Pinto, Meenakshi Chakraborty, Antonina Andreeva, Gabriele Ghiotto, Jim Shaw, Amy D. Proal, Justin L. Sonnenburg, Ami S. Bhatt, Simon Roux, Georgios A. Pavlopoulos, Stephen Nayfach, Nikos C. Kyrpides

## Abstract

Viruses are key modulators of human gut microbiome composition and function. While metagenomic sequencing has enabled culture-independent discovery of gut bacteriophage diversity, existing genomic catalogues suffer from limited geographic representation, sparse taxonomic classification, and insufficient functional annotation, hindering detailed investigation into phage biology. Here, we present the Unified Human Gastrointestinal Virome (UHGV), a collection of 873,994 viral genomes from globally diverse populations that addresses these limitations. UHGV provides high-quality virome references with extensive host predictions, comprehensive functional annotations, protein structures, a classification framework for comparative analysis, and a web portal to facilitate data access. Using UHGV to profile worldwide metagenomes, we found that host range breadth is strongly associated with phage prevalence. Additionally, we identified diversity-generating retroelements and DNA methyltransferases as key factors enabling phage populations to access diverse hosts, revealing how specific genomic features contribute to global phage distribution patterns. UHGV is available at http://uhgv.jgi.doe.gov.

## Introduction

The human gut microbiome is a complex ecosystem that harbours a vast diversity of microorganisms, profoundly influencing human health by modulating immune responses, nutrient metabolism, and disease susceptibility^1^. The viral components of these communities, dominated by bacteriophages (phages) that infect bacteria, play important roles in shaping microbiome structure and function in both health and disease through predation and horizontal gene transfer. This phage-mediated modulation of gut bacteria influences metabolic outputs and pathogen proliferation^2^, and can induce virulence in several bacterial species^3^, thereby impacting human health. For instance, shifts in the composition of the virome have been linked to various conditions, including inflammatory bowel disease^4^, metabolic syndrome^5^, and obesity^6^. Moreover, there is growing interest in harnessing phages and their proteins as alternatives to traditional antibiotics for treating infections, through methods such as phage therapy^7^ and endolysin-based therapy^8^.

In recent years, analysis of metagenomic sequencing data has revolutionised our ability to survey the gut microbiome and its uncultivated viruses, enabling culture-independent recovery of microbial and viral genomes directly from faecal samples. The rising availability of gut metagenomes, produced from bulk microbial communities or samples enriched for virus-like particles, combined with advances in bioinformatics methods, has yielded valuable insights into viral diversity, host interactions, and biogeographical distribution. These efforts have led to the discovery of major viral clades, such as the *Crassvirales*, the most prevalent gut phages in Western adults^9^, and the *Grandevirales* phages, whose genomes can exceed 500 kb^10^. To comprehensively characterise gut viral diversity, multiple studies have performed systematic identification of virus genomes across large collections of human gut metagenomes, producing catalogues that encompass diverse viral lineages^11– 24^.

Despite significant progress in cataloguing gut viral diversity, virome research faces multiple longstanding challenges. First, most available human gut virus catalogues fail to serve as comprehensive references because they either target specific populations or are biased towards Western populations while underrepresenting non-industrialised societies. Second, the taxonomy maintained by the International Committee on Taxonomy of Viruses (ICTV) has limited coverage of uncultivated viruses, leaving most gut phages unclassified^25^ and hindering cross-study interoperability, as researchers often resort to ad hoc comparative genomics methods. Lastly, several existing catalogues predate important methodological advances in virus identification and host prediction, as well as breakthroughs in protein annotation and structure prediction. These latter developments could be leveraged to address the longstanding challenge of functional annotation of viral proteins, which limits mechanistic insights into virus biology.

To address these critical deficiencies, we developed the Unified Human Gastrointestinal Virome (UHGV), the most comprehensive gut virome resource to date. UHGV was generated through rigorous integration and quality control of multiple existing catalogues and comprises 873,994 genomes and 37,443,649 proteins from uncultivated human gut viruses. This resource overcomes existing limitations by incorporating virus genomes from diverse global populations, implementing a novel taxonomy-like classification framework to systematically organise viral diversity, and providing deep functional annotations complemented by predicted protein structures. In addition to these core features, UHGV includes extensive supporting data, such as biogeographical profiles and single nucleotide variant analyses, to support a broad range of research applications. We anticipate that UHGV will accelerate human gut virome research across all scales, from individual genes to global distribution patterns.

## Results and Discussion

### Creating a curated, comprehensive, and annotated catalogue of human gut viruses

To establish a comprehensive catalogue of viruses from human gastrointestinal microbiomes, we collected 2,242,702 putative uncultivated virus genomes from 12 published studies^11–19,22– 24^. To ensure reliability and uniformity across these heterogeneous catalogues, we developed a quality control procedure incorporating recent advances in virus genome identification and quality assessment (Figure 1A). Through this procedure, we were able to remove sequences with insufficient evidence of viral origin (61.0% of the initial dataset). As short genome fragments cannot always be unambiguously identified as viral, we partitioned the retained genomes into confident (88.2%) and uncertain (11.8%) classifications based on the strength of supporting evidence. Subsequently, all genomes were trimmed to remove host genes flanking integrated viruses (proviruses), and stratified into quality tiers based on their estimated completeness (Figure S1A, Table S1). This rigorous quality control resulted in an improved quality over prior resources and was critical to ensure the reliability of the data to draw biological insights. For example, we removed several sequences that, although highly prevalent in metagenomes, lacked viral genes and showed hallmarks of mobile genetic elements rather than viruses. One notable case, a genomic island containing multiple integrated elements, ranked among the top 0.73% most prevalent sequences across bulk metagenomes but contained no identifiable viral genes (Figure S1B).

**Figure 1.**
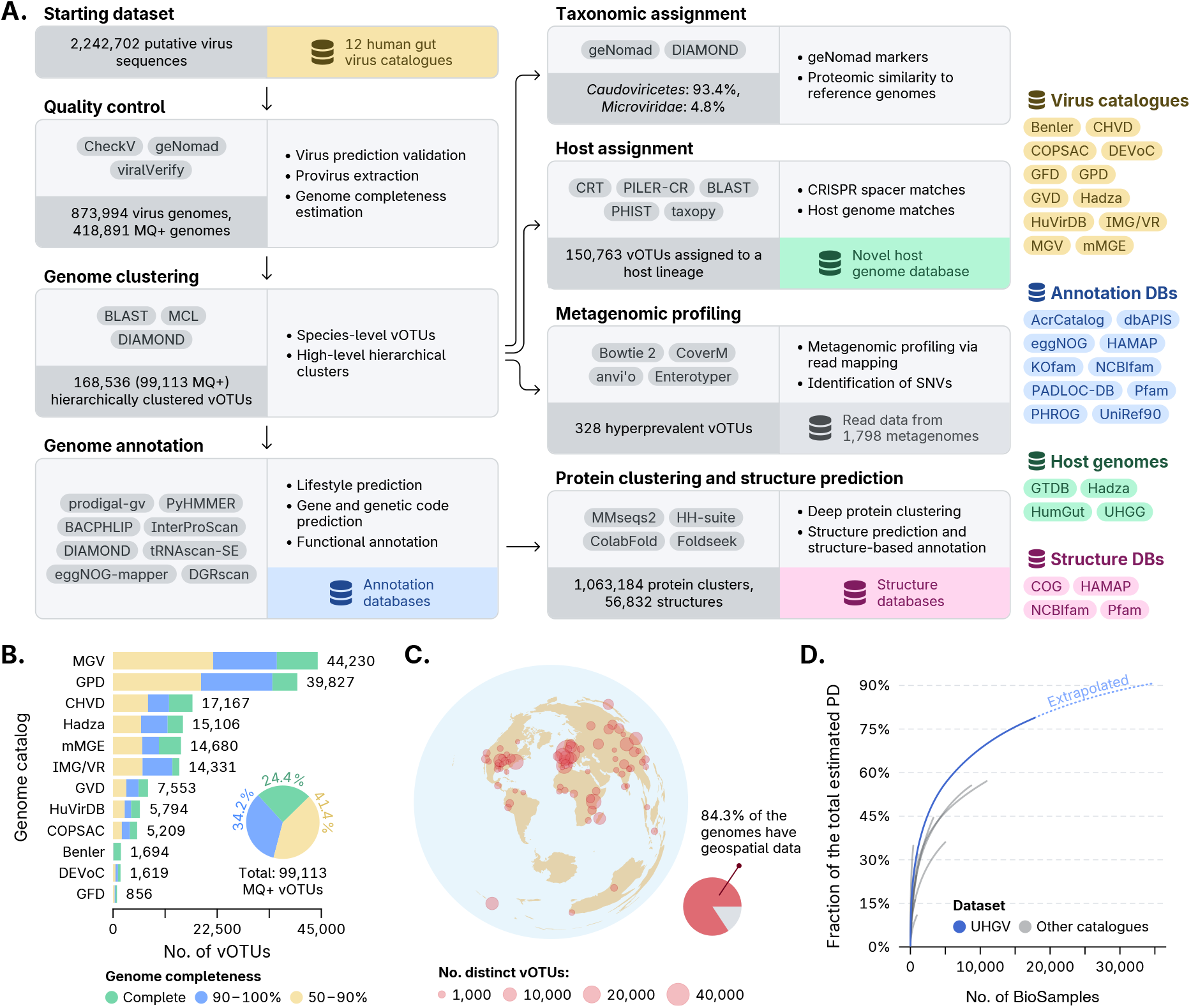
**(A)** Schematic of the pipeline for constructing the UHGV dataset. **(B)** Bar chart displaying the number of vOTUs (x-axis) among the medium-quality or better UHGV genomes with confident virus prediction from each source catalogue (y-axis). Bar colours indicate the fractions of vOTUs within each quality tier. The pie chart displays the overall proportion of each quality tier across all catalogues. **(C)** Geographical distribution of UHGV genomes with available geospatial data. Circle sizes on the map are proportional to the number of distinct vOTUs in each region. For visual clarity, samples from nearby locations were clustered using DBSCAN, and the coordinates of the cluster medoids were plotted. Samples lacking specific coordinates but containing country-level metadata were positioned at the centroid of their respective countries. The map is presented using the azimuthal equidistant projection. **(D)** Accumulation curve depicting relative Faith’s phylogenetic diversity (y-axis) as a function of the number of BioSamples (x-axis) where phage genomes were identified. The blue curve represents the UHGV dataset, and the grey curves correspond to its constituent datasets. Diversity was computed from medium-quality or better vOTU representatives and is expressed relative to the total expected diversity, accounting for the estimated unobserved diversity. The dashed blue line extrapolates the curve, indicating the anticipated rate of discovery for unobserved diversity. PD: phylogenetic diversity.

The resulting UHGV catalogue comprises 873,994 viral genomes, representing 837,790 unique sequences that were grouped into 168,536 virus operational taxonomic units (vOTUs; species-level groups of genomes with ≥95% nucleotide identity over ≥85% of genome length)^26^. Of these, 99,113 vOTUs contain at least one genome with confident virus identification and of medium-quality or better (genome completeness ≥50%), more than doubling the size of the largest prior catalogue (Figures 1B and S1A). Notably, 72.1% of UHGV genomes belong to vOTUs containing high-quality (genome completeness ≥90%) or complete representative genomes, a significant improvement over existing resources (Table S1). This high completeness of vOTU representative genomes (the highest-quality genome in the vOTU) is critical for overcoming assembly fragmentation challenges in metagenomic data, as such genomes can be used as references. UHGV’s robust foundation for the study of human gut viruses is further strengthened by its broad coverage, with genomes sourced from 19,008 identifiable sequencing samples deposited over 14 years (Figure S1C) and collected from individuals across 42 countries (Figure 1C, Table S1), representing diverse human populations. Reflecting its extensive size and breadth, UHGV markedly expands the known phylogenetic diversity^27^ of gut viruses, achieving a 1.5-fold increase over the previous largest catalogue^13^ and capturing 78.9% of the estimated total phylogenetic diversity of gut *Caudoviricetes*, the dominant class of double-stranded DNA phages (Figures 1D and S1D, Table S2).

To produce a reliable and extensive data resource to support virus research, we designed a pipeline (Figure 1A, see Methods for details) that thoroughly annotates UHGV genomes, including lifestyle classification (lytic or temperate), prediction of protein-coding genes and tRNAs, and protein function annotation using a wide range of reference databases. Protein sequences were also subjected to deep clustering, structure prediction, and structure-based functional annotation. To organise the virus genome diversity within an established framework, we assigned 99.98% of the vOTUs to an ICTV taxa^28^, though only 16.8% were classified at or below the family rank (Table S3). Recognising the importance of virushost interactions in understanding viral biology, we combined multiple host prediction methods with a new database of human gut prokaryote genomes to assign 89.5% of vOTU representatives to putative host taxa (Table S3). In addition, we mapped reads from 1,798 human gut metagenomes — representing populations spanning varied geographies and lifestyles — to vOTU representative genomes, enabling analysis of global virus distribution and investigation of viral diversity at singlenucleotide resolution.

### Hierarchically classified high-completeness genomes capture most of the intestinal virome diversity

To investigate *Caudoviricetes* diversity within UHGV, we reconstructed a comprehensive phylogeny based on concatenated marker genes from 417,277 genomes, representing 84,805 vOTUs and encompassing 81.3% of the high-quality and complete vOTU representative genomes in the dataset. Clustering the phylogeny into 951 clades revealed that UHGV includes at least one complete genome for most of these lineages (green outer ring in Figure 2A). Through the evaluation of phylogenetic diversity, we found that phage diversity is distributed across countries with diverse populations (Figure 2B), and that no single source catalogue accounts for more than 65% of the total diversity in UHGV (Figure 2C), highlighting the importance of a unified resource to comprehensively capture global phage diversity.

**Figure 2.**
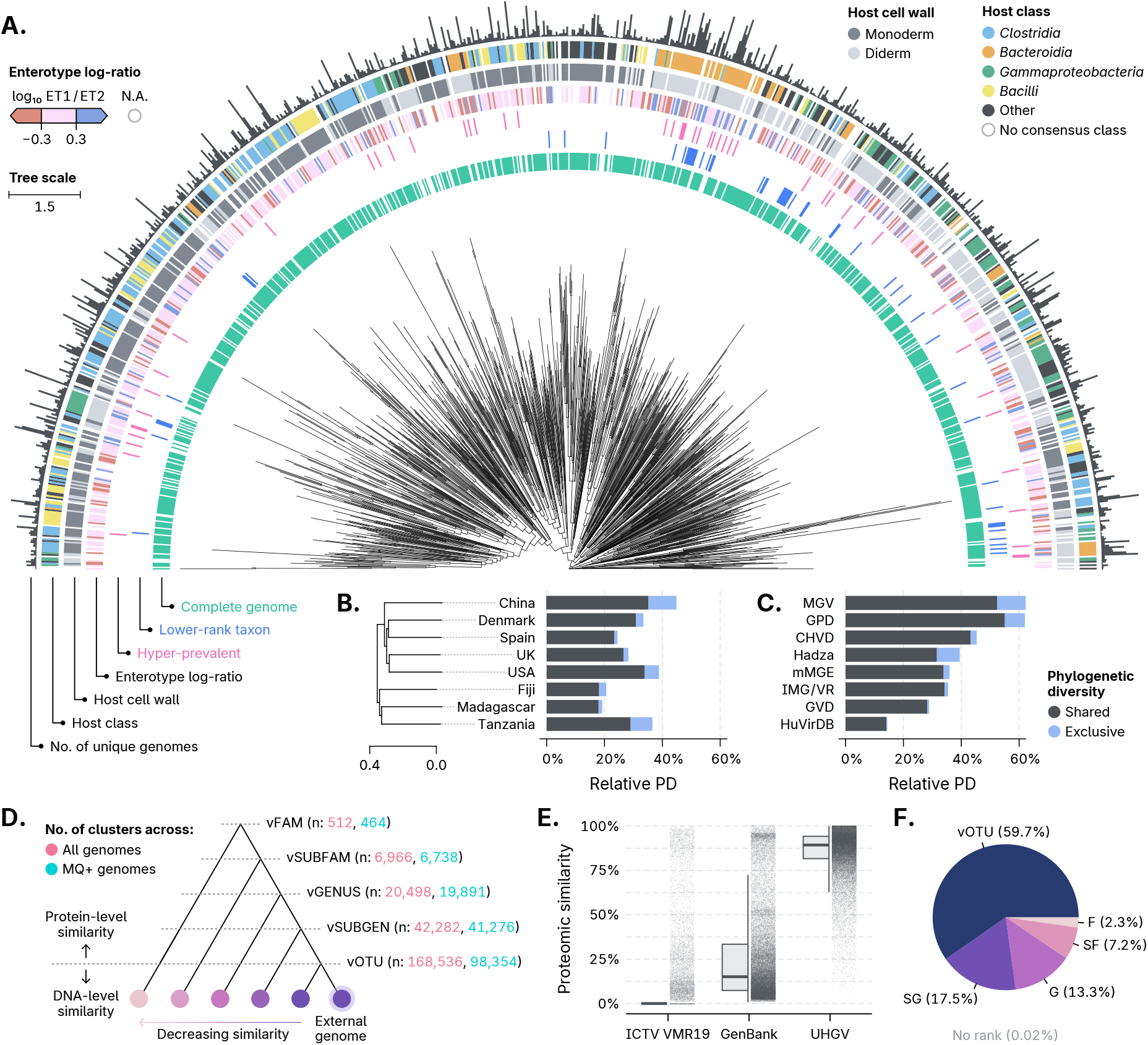
**(A)** Phylogenetic tree of *Caudoviricetes* phages from UHGV. For visual clarity, low-quality genomes were excluded, and vSUBFAM representative genomes were clustered by cophenetic distances using DBSCAN, resulting in the 951 displayed tips. Clades with support <70% were collapsed. Outer rings display metadata for the genomes represented by each tip. **(B**,**C)** Bar charts representing the relative Faith’s phylogenetic diversity of *Caudoviricetes* (x-axis) found within the most diverse **(B)** countries of origin and **(C)** source catalogues. Blue portions indicate diversity exclusive to each catalogue or country. Countries are arranged in a BIONJ dendrogram, based on UniFrac distances computed from rarefied samples (≤500 genomes/country). **(D)** UHGV genomes are organised in a taxonomy-like cluster hierarchy, starting with species-level vOTUs based on DNA similarity and progressing to broader ranks determined through protein similarity. External genomes are classified based on their similarity to UHGV genomes. **(E)** Distribution of proteomic similarity scores (y-axis) for 44,290 phage genomes from recent studies, compared against three reference databases (x-axis). For each queried phage, only the highest similarity score within each database is shown. **(F)** Pie charts showing the taxonomic classification specificity of the 44,290 phage genomes assigned to UHGV hierarchical clusters. Each genome was classified to the most specific rank possible. The percentage of genomes not assigned to any UHGV cluster is shown at the bottom. ET1: enterotype 1; ET2: enterotype 2; PD: phylogenetic diversity; MQ+: genomes of medium-quality or better; SG: vSUBGEN; G: vGENUS; SF: vSUBFAM, F: vFAM.

A major challenge in viral metagenomics is that most uncultivated phages do not belong to existing named taxa at low ranks (i.e., below the class level)^29^. In UHGV, only 11.0% of the *Caudoviricetes* vOTUs — representing 7.7% of the total phylogenetic diversity — were assigned to a taxon at or below the order rank (blue outer ring in Figure 2A). This taxonomic paucity poses a major barrier to robust comparative genomics and fine-grained metagenomic profiling analyses, compelling researchers to use varied bespoke methods that undermine cross-study comparisons. To address these challenges, we developed a taxonomy-like framework to organise all UHGV vOTUs into hierarchical clusters based on proteomic similarity (Figure 2D). The resulting ranks (named vSUBGEN, vGENUS, vSUBFAM, and vFAM) roughly correspond to the ICTV’s subgenus, genus, subfamily, and family classifications, allowing our framework to outperform a state-of-the-art method^30^ in reconstructing ICTV taxa from genome data alone (Table S4). To enable adoption of this framework in future studies, we developed a toolkit to consistently assign external viral genomes to UHGV clusters.

Discretizing phage diversity into ranks of varying resolution enables assessment of human gut phage diversity saturation at different levels. This reveals that, although diversity at the vOTU level is far from saturation (open accumulation curves, γ: 0.73), vFAM-level diversity approaches exhaustion (γ: 0.13) (Figure S2A). Cross-country comparisons further demonstrate that understudied populations harbour substantial unexplored viral diversity (Figure S2B), even at higher ranks, and that sampling of new countries increases diversity faster than random sampling (Figure S2C), emphasising the need for broader geographic sampling.

To further evaluate the extent of the human gut phage diversity captured by UHGV, we assessed its coverage of the diversity among 44,290 phage genomes from metagenomes of diverse human gut microbiomes from four studies^20,21,31,32^ published after UHGV was compiled. After reprocessing these data using a consistent pipeline, we found that the identified phage genomes were highly similar to those in UHGV (median: 89.1% proteomic similarity), while exhibiting little similarity to ICTV species exemplars and GenBank viruses (Figure 2E, medians: 0.0% and 15.1%). As a result, 99.98% of the genomes from the selected studies were assigned to UHGV genome clusters, most at the vOTU level (Figures 2F and S3A). Among genomes assigned to vOTUs, the UHGV vOTU representatives were generally more complete than the external genomes matching them (Figure S3B). Consistent with its extensive coverage, UHGV outperforms existing resources in gut virome profiling, producing higher mapping rates for infant gut datasets (Figure S3C, Table S5, Supplementary Note 1). These results underscore that UHGV captures much of the gut viral diversity and offers high-quality reference genomes for most viral groups in the human gut microbiome.

### Deep annotation of the protein landscape of human gut viruses

The analysis of viral proteins is essential for extracting biological insights from genomic data, yet inferring protein function from sequence alone remains difficult as most viral proteins lack functional annotation^33^. This stems from the vast and rapidly evolving virus protein repertoire, which limits traditional sequence similarity-based annotation methods. This rapid diversification also obscures homology relationships, hindering the study of virus evolution. To address these challenges and enable deeper understanding of viral function and evolution through UHGV, we set out to determine homology relationships across the UHGV proteome and perform exhaustive functional annotation of these proteins.

To efficiently group divergent homologs, we developed a two-step clustering strategy designed to enhance sensitivity (Figure 3A). First, we performed all-against-all protein alignments to generate intermediary groupings, which were then used in a more sensitive profile-versus-protein search and a second round of clustering. Using this approach, the full set of 37,443,649 UHGV proteins was grouped into 1,063,184 clusters, of which 474,134 contain multiple distinct sequences (Figure S4A). Next, protein structures were predicted for representative proteins from clusters with at least 15 unique sequences, yielding 56,832 high-quality structures (median avg. pLDDT: 81.4; 79.5% with avg. pLDDT ≥70) representing 70.1% of the UHGV protein repertoire (Figure S4B, Table S6). These structures show substantial novelty, as 90.3% lack structurally redundant matches (≥90% coverage) in PDB, and 37.3% have no redundant matches in AlphaFold DB^34^.

**Figure 3.**
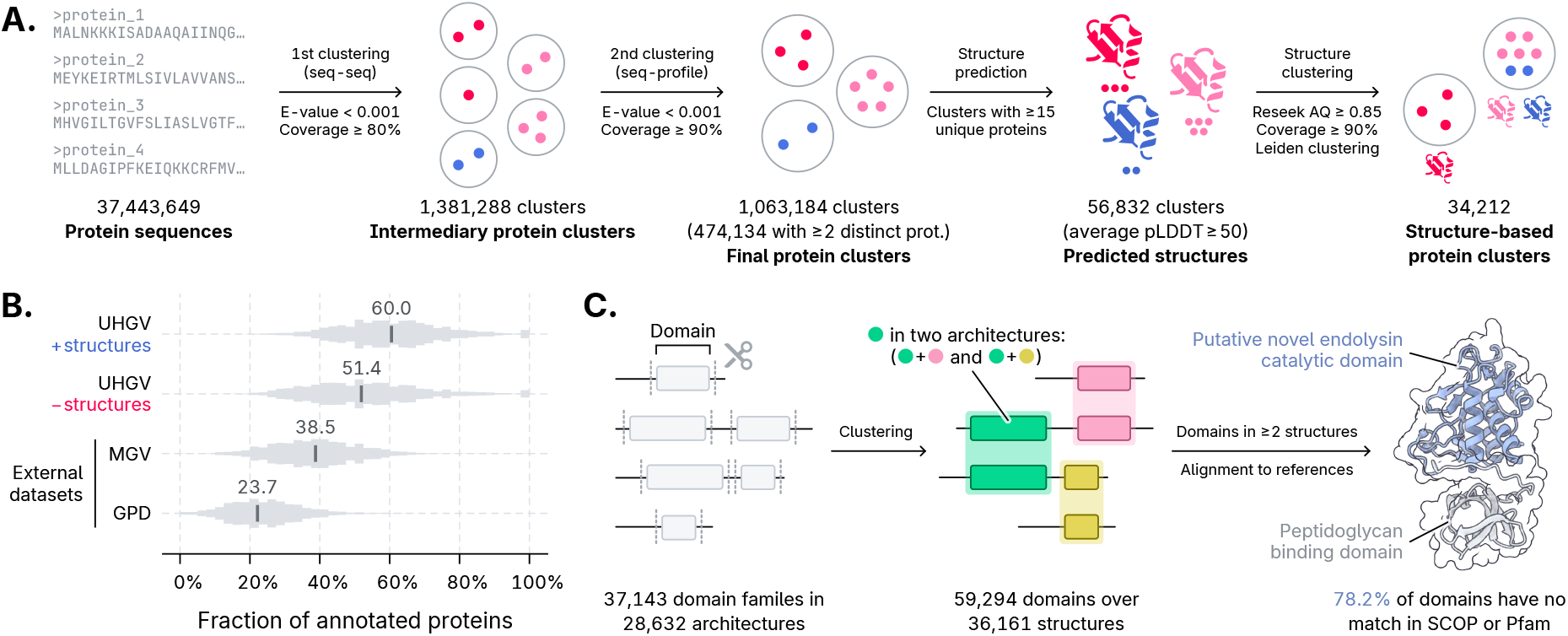
**(A)** Pipeline for two-step protein clustering, structure prediction, and structure clustering. **(B)** Distributions of the protein annotation rate across genomes within the GPD, MGV, and UHGV datasets. For UHGV, two distributions are shown: with (top) and without (bottom) structure-based annotation. **(C)** Pipeline for identifying novel domains in UHGV protein structures. Following automated domain segmentation, high-confidence domains were clustered, and protein domain architectures were determined. Domains detected in multiple structures were aligned to SCOP and predicted structures of Pfam domains to assess novelty. A likely novel endolysin catalytic domain (highlighted in blue) is shown on the right.

Protein structure is more conserved than sequence^35^, allowing structure-level alignments to detect remote homologies missed by sequence-based methods. To identify such remote relationships within UHGV, we grouped proteins based on structural similarity among representative models, producing 34,212 structure-based clusters. Among these clusters, the largest contained 141,087 distinct orthologues of the terminase, a protein that, despite being nearly universal in *Caudoviricetes*, did not appear among the largest sequence-based clusters due to high sequence divergence (Figure S4C– E). More broadly, analysis of accumulation curves of both sequence- and structure-based clusters showed that UHGV captures the majority of protein diversity in the human gut virome, with near-complete saturation at the structure level (Figure S4F).

The challenge of limited viral protein functional annotation was addressed through two approaches: integration of comprehensive annotation resources and structure-based annotation. Proteins from medium-quality or better vOTU representatives were annotated via alignment to reference proteins and HMMs, using 11 tools and databases covering diverse functions. This resulted in a higher annotation rate in UHGV compared to other catalogues, with a median of 51.4% annotated proteins per genome, versus 23.7% in GPD and 38.5% in MGV (Figure 3B, Table S7). To further improve annotations, UHGV protein structures were aligned to newly predicted structures from four reference annotation databases (Figure S4B), increasing the median annotation rate to 60.0% (Figure 3B, Table S7). A benchmark comparing structure-based annotations to HMM-based Pfam annotations showed strong concordance at the family (81.3%) and clan (95.3%) levels (Figure S4G), supporting the reliability of the structure-based annotation.

The availability of a large set of viral protein structures also enables domain-level evolutionary analysis, offering deeper insight into viral protein function and evolution, as domain shuffling plays a major role in driving functional innovation in phages^36^. Through domain segmentation of protein structures and structure-based clustering of individual domains, we identified 37,143 domain families occurring in multiple structures (Figure 3C). Among these, 74.7% are detected in multiple domain architectures, with 13.4% found in five or more distinct architectures, highlighting extensive domain shuffling within the UHGV proteome. Alignment of domains found in at least two structures against SCOPe and Pfam revealed that 78.2% lack significant matches. Detailed examination of one such domain, which co-occurs with several lysis-related domains, revealed that it likely is a novel papain-like catalytic domain of endolysins (Figure S5, Table S8, Supplementary Note 2), demonstrating the power of this approach for discovering novel viral protein functions.

### Worldwide gut virome profiling identifies host range breadth as a driver of prevalence

We examined the geographic origins of UHGV genomes and found a significant association between geographic distance and virus community composition (Mantel *r*: 0.3882, *p* value: 0.006; Figures 2B and S6A,B), indicating that phage communities are differentiated among human populations from different geographic locations. To investigate these biogeographic patterns in greater detail, we profiled 1,798 bulk metagenomes and 673 viromes — selected to represent diverse human populations with varied diets — to detect and quantify both viral and prokaryotic components across cohorts with different nutritional regimes worldwide. As phage community makeup is shaped by the prokaryotic component, we evaluated whether human gut phage communities are structured, forming distinct clusters that parallel the enterotypes described for bacterial gut communities^37^. Through *de novo* clustering, we found that viral components form two major clusters that roughly correspond to enterotypes 1 (ET1) and 2 (ET2), though with weaker structure (Figures 4A,B and S6C). Importantly, these clusters reflect human dietary differences: the ET1-like samples were majorly from industrialised populations, while ET2-like samples were mostly from hunter-gatherers and transitioning societies (Figure S6D,E).

**Figure 4.**
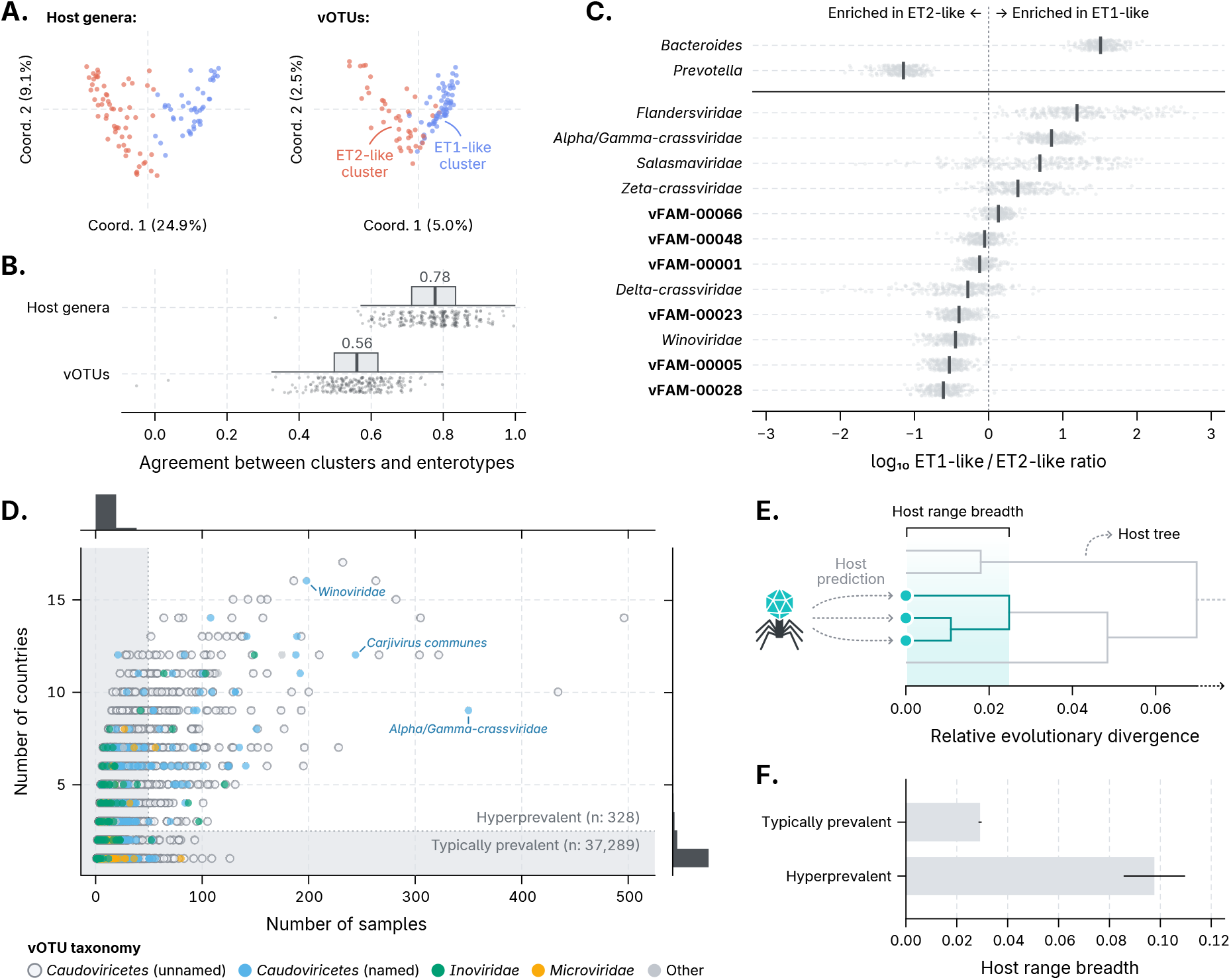
**(A)** Multidimensional scaling ordinations of 105 randomly selected human gut communities, colour-coded by k-medoids clusters based on Jensen-Shannon distances. Blue and red indicate clusters resembling enterotype 1 (ET1-like) and enterotype 2 (ET2-like), respectively. Left: host genus abundances; right: vOTU abundances. Percentages on the axes labels indicate the proportion of variance explained by each axis. **(B)** Agreement between host-(top) and vOTU-based (bottom) clustering and reference enterotype classification. Each circle represents a cluster from one of 250 iterations using random community subsets (≥15 countries, ≤40 samples per country). Box plots show adjusted Rand index values quantifying agreement between inferred clusters and enterotype labels, with medians shown above each box. **(C)** Association of the 12 most abundant vFAMs with enterotype 1-like (ET1-like) and 2-like (ET2-like) community clusters, expressed as the log ratio of the mean relative abundance in ET1-like versus ET2-like samples. Circles represent values from 250 iterations; vertical lines indicate medians. Reference distributions for *Bacteroides* and *Prevotella* are shown at the top. **(D)** Frequency of vOTU detection (circles) across samples (x-axis) and countries (y-axis), colour-coded by taxonomy. The shaded area indicates vOTUs not meeting hyperprevalence criteria. Marginal histograms show vOTU distribution along each axis. **(E)** The host range breadth of each vOTU was quantified from the phylogenetic distances among the host genomes to which it was linked. **(F)** Mean host range breadth (x-axis) for typically prevalent versus hyperprevalent vOTUs. Horizontal lines represent the standard error of the mean.

Recent studies have identified certain phage groups, such as *Crassvirales*, as dominant in the human gut^18,38^, however these investigations were biased toward well-studied populations. Given our findings on phage community differentiation across enterotypes, we asked whether ET2-like communities harbour highly abundant phage groups not previously recognised as such. Among the 12 most prevalent vFAMs in balanced sample subsets (to mitigate bias from country of origin or diet), 71.4% of those enriched in ET2-like communities lacked assignment to a known taxon at the order level or below, compared to only 20.0% in ET1-like (Figure 4C). Next, we evaluated individual vOTUs in terms of prevalence, defining hyperprevalent vOTUs as those detected in ≥50 samples from ≥3 countries (Figure 4D). Although only 328 (0.9%) vOTUs met this criterion, they were present in 92.5% of the evaluated datasets and accounted for 18.6% of all detections. Despite their broad distribution, the majority (83.2%) remain unclassified at the family level, and among the fraction that are classified, phage clades previously described as highly abundant were generally depleted in ET2-like communities (Figure S6F,G).

While the exceptional prevalence of a few phages is recognised, the mechanisms underlying the variations in prevalence are unclear. The prevalence of bacterial taxa does not reliably predict that of associated phages, which can vary widely within the same host taxon (Table S9). This decoupling suggests that phage-specific traits, not just host factors, shape their distribution. Therefore, we hypothesised that phages with greater host-switching ability are more likely to be highly prevalent. To test this, we devised a new metric to quantify the host range breadth (HRB) of a phage population based on the phylogenetic distance among host taxa linked to each vOTU (Figures 4E and S6H). Using this approach, we found a strong positive link between HRB and prevalence, with hyperprevalent vOTUs exhibiting significantly higher HRB than typically prevalent ones (Mann-Whitney *U p* value < 0.001; Figures 4F and S6I). This relationship aligns with a scenario where enhanced host-switching potential contributes, alongside other factors, to higher phage prevalence by facilitating access to diverse ecological niches and, consequently, resilience to host population fluctuations.

### Identification of genetic factors enabling broad host range in phages

The newly devised HRB metric enabled quantitative assessment of host range across phages. In UHGV, 86.7% of vOTUs exhibited a species-level host range, with most of the remainder being either genus- or family-level (7.4% and 4.7%, respectively; Figure S7A). Examination of HRB patterns revealed marked differences between *Caudoviricetes* and ssDNA phages, as well as differences tied to genome length, host classes, and lifestyles (Figure S7B). Notably, HRB showed a significant phylogenetic signal within *Caudoviricetes* (Pagel’s λ: 0.97, *p* value < 0.001), indicating that closely related phages tend to exhibit similar HRB and implying that genetic factors are key determinants of this trait. Accordingly, gene annotations present in ≥1% of complete *Caudoviricetes* genomes explain 47.4% ± 0.5% of their HRB variance.

Motivated by these findings, we screened for protein families and domains linked to increased HRB, uncovering 65 functions with significant positive associations (*t* test FDR < 0.05; Figure 5A, Table S10). Among the strongest associations, three major biological processes stood out: diversity-generating retroelements (DGRs), counter-defense systems — especially DNA methyltransferases (MTases) —, and cell wall lysis. Interestingly, these three processes operate at different stages of the phage infection cycle, as DGRs influence phage adsorption, counter-defence systems enable evasion of host immunity, and endolysins degrade the cell wall to induce host lysis and release progeny (Figure 5B). The association between these processes and elevated HRB remained consistent across alternative statistical methods and HRB quantification approaches (Table S10).

**Figure 5.**
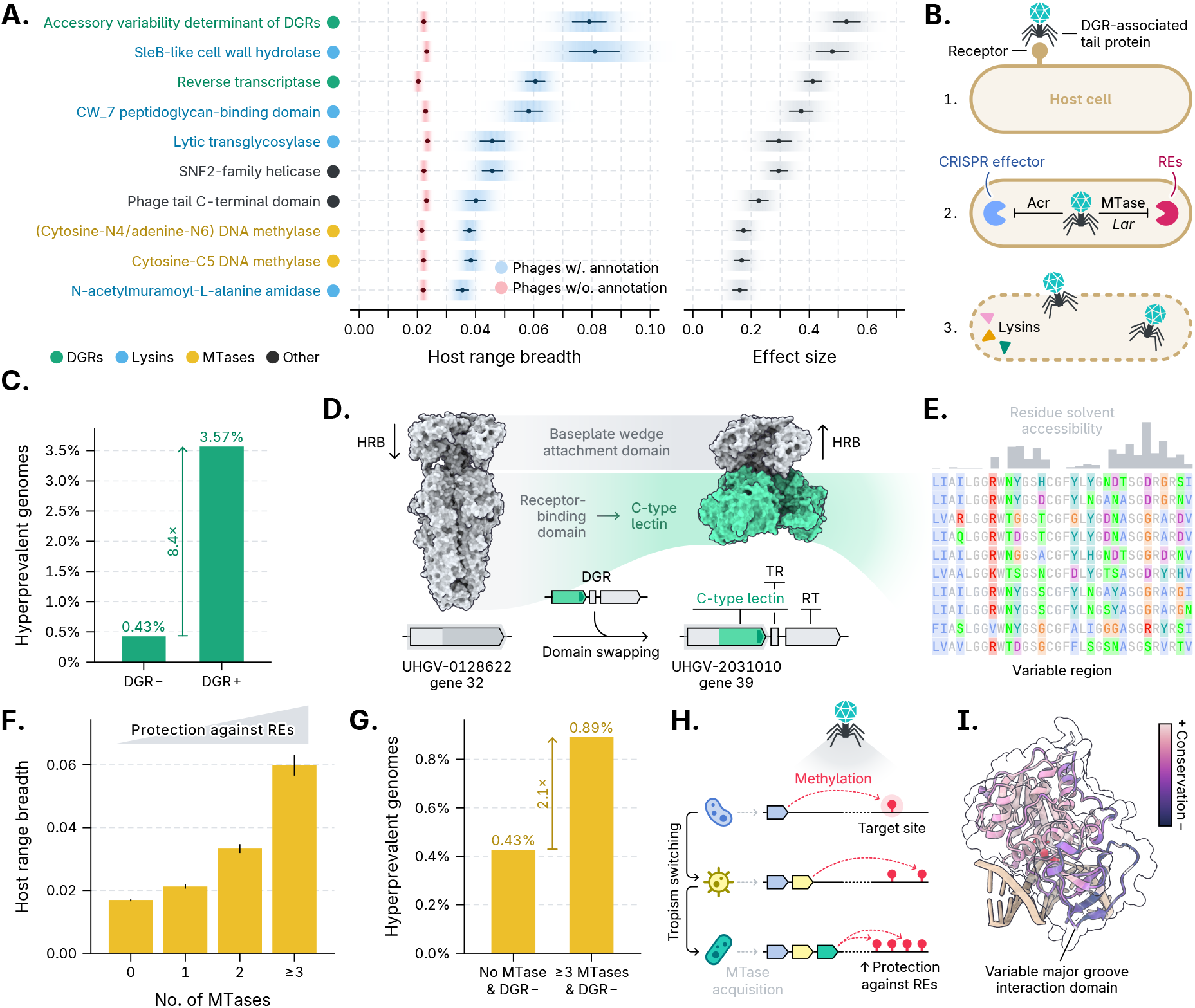
**(A)** Top ten functional annotations most strongly associated with increased host range breadth (HRB). Left: average HRB (x-axis) of phages encoding (blue) or lacking (red) each function. Right: effect size (Cohen’s *d*, x-axis) on HRB. Horizontal lines represent 95% bootstrap confidence intervals. Function names are colour-coded by biological process. **(B)** Functions linked to increased HRB map to three events of the infection cycle: (1) adsorption, (2) counter-defence, and (3) lysis. **(C)** Proportion of hyperprevalent genomes (y-axis) in phages with or without a diversity-generating retroelement (DGR+ and DGR−, respectively). **(D)** DGR acquisition typically involves domain swapping, replacing an existing receptor-binding domain with a DGR-targeted C-type lectin domain. **(E)** Alignment of the variable region of gene 39 from the UHGV-2031010 genome across five metagenomes. Polymorphic residues are coloured. The bar plot above shows the relative solvent accessibility of each residue. **(F)** Mean HRB (y-axis) increases with DNA methyltransferase (MTase) copy number (x-axis) among DGR− phages. **(G)** Proportion of hyperprevalent genomes (y-axis) in DGR− phages lacking MTases versus those encoding three or more MTases. **(H)** MTases are frequently acquired via horizontal gene transfer and may accumulate through successive infections of different hosts, thus increasing genome methylation and protection against restriction enzymes. **(I)** Relative evolutionary rates of UHGV C5C-MTase residues mapped onto the HhaI structure39 (PDB: 1MHT), with darker colours indicating faster-evolving sites. RE: restriction enzyme; TR: template region; RT: reverse transcriptase.

DGRs were the strongest genetic determinant of high HRB and, in turn, of phage prevalence, as evidenced by *Caudoviricetes* encoding this system being 8.4-fold more likely to be hyperprevalent and accounting for 50.9% of hyperprevalent vOTUs (Figure 5C, Table S11). DGRs operate through an error-prone reverse transcriptase that introduces mutations in a target variable region of a target C-type lectin domain, thereby diversifying its receptor-recognition surface and altering its binding specificity (Figure S8A)^40^. Within UHGV, DGRs occur in phages infecting a wide range of host taxa and typically target tail proteins (Figure S8B,C), which mediate host recognition. Structural analysis of these DGR-targeted proteins revealed pronounced diversity (Figure S8D), with C-type lectin domains exhibiting a 6.8-fold higher rate of domain shuffling than average, consistent with extensive recombination and frequent horizontal transfer^41^. To investigate DGR acquisition by phages in detail, we examined pairs of phages from the same vFAM that differed in DGR presence. This analysis revealed events where DGR acquisition, coupled with an increase in HRB, involved domain swapping in tail proteins, replacing the original host-binding domain with a C-type lectin domain while retaining the portion that interacts with the rest of the virion (Figures 5D and S8E,F). We also assessed the impact of DGRs on tail protein diversification by analysing genomic variation and selection pressures at single nucleotide resolution through read mapping data (Supplementary Note 3). This revealed that natural selection favours diversification of surface-exposed residues in the variable region of DGR targets (pN/pS^42^: 2.9; Figure S8G–J), promoting missense mutations that often replace amino acids with others of distinct physico-chemical properties (Figure 5E), likely affecting host tropism.

Phage infection requires not only cell entry but also successful completion of the replication cycle, a process that depends on additional phage-encoded functions. Accordingly, the association between HRB and prevalence remains significant even when controlling for DGRs (Figure S9A). Among functions associated with increased HRB beyond DGRs, several involve evading host immunity (Table S10), with MTases, which protect phage genomes from host restriction enzymes (REs) via DNA methylation, being prominent^43^. Because bacterial RE repertoires vary, we reasoned that phages with multiple MTases, and thus broader RE protection, would have greater chance of success across diverse hosts^44^. Supporting this hypothesis, MTase count strongly correlated with increased host range breadth (Figure 5F) while controlling for DGRs, genome length, and lifestyle (phylogenetic regression *p* value < 0.001; Figure S9B–D). Furthermore, among phages without DGRs, those encoding ≥3 MTases were 2.1 times more likely to be hyperprevalent than those lacking them (Figure 5G, Table S12). To investigate how phages accumulate MTases, we analysed the phylogenetic distribution of these enzymes across *Caudoviricetes* and found them to be highly homoplasic (Figure S9E), indicating frequent horizontal transfer. Such acquisitions may occur sequentially as phages infect different hosts (Figure 5H), with each event introducing MTases with distinct sequence specificities, as suggested by the high variation within their DNA-binding domains (Figures 5I and S9F,G). The resulting accumulation would progressively increase DNA methylation, enhancing virus protection against host REs in subsequent host transitions. Examination of factors that might affect MTase acquisition revealed that temperate phages are more likely than lytic ones to accumulate MTases (Figure S9H). In addition, phages with a greater number of MTases were more likely to carry an HU DNA-binding protein, with temperate phages encoding ≥4 MTases showing a 19.0-fold enrichment for HU compared to those lacking MTases (phylogenetic regression *p* value < 0.001; Table S13). As HU has been shown to bend DNA and promote recombination^45^, these findings are consistent with recombination rates — influenced by factors such as lifestyle and HU — affecting MTase acquisition and host range breadth.

Determining how specific endolysin domains are associated with HRB is challenging, as it depends on factors such as the number and type of endolysins encoded by a given phage, the combined effects of enzymatically active domains (EADs) and carbohydrate-binding domains (CBDs), and the composition of the host cell wall. Despite this complexity, we found that the EADs and CBDs linked to increased HRB target conserved bacterial cell wall structures, whereas the only two evaluated EADs not associated with elevated HRB are selective for specific peptide stem variations^46,47^ (Table S14). This is consistent with a model in which generalist endolysin domains offer greater adaptive value to phages transitioning to new hosts^48^. To assess endolysin diversity across hosts, we leveraged UHGV’s extensive set of 68,920 unique endolysins (Figure S10A–C, Table S15, Supplementary Note 4) contextualised by phage–host predictions. We found that bacterial cell wall architecture is a strong determinant of endolysin domain architecture, with certain CBDs and EADs exhibiting differential distribution across host clades (Figure S10D,E, Table S16). Finally, we examined sequence diversity in two prevalent CBDs that bind conserved cell wall structures. Both showed strong co-evolution with their hosts (Figures S10F–I), indicating adaptation to subtle, taxon-specific cell wall differences and illustrates the evolutionary interplay between phage proteins that directly interact with the host and the hosts themselves.

## Conclusion

To support human gut virome research and address key limitations in the field, we present UHGV, a genomic resource that integrates multiple virome datasets into a single highquality reference. The viral genomes in UHGV are thoroughly annotated to support diverse investigations into virus biology, incorporating data spanning from worldwide biogeography to nucleotide-level sequence variation. Leveraging this data, we investigated global patterns of phage occurrence and demonstrated that phage prevalence is strongly shaped by host range breadth. Building on this finding, we developed a mechanistic framework to explore the genetic factors underlying phagehost interactions, enabling us to identify specific mechanisms through which functions such as DGRs and MTases facilitate phage transitions between different hosts. These results illustrate how data-driven approaches, empowered by UHGV, can yield insights into phage biology. Beyond fundamental virology, these insights have practical applications for phage engineering and the development of phage-based and lysin-based therapies. Moving forward, experimental verification of the proposed host range expansion mechanisms is imperative, alongside the development of more sophisticated computational methods for predicting phage-host specificity and infection outcomes.

We anticipate that the UHGV will become an essential resource for studying viruses in the human gut. To facilitate its adoption, we provide a user-friendly website for data exploration and download (http://uhgv.jgi.doe.gov/), as well as a command-line toolkit for assigning phage genomes to clusters within the UHGV taxonomy-like genome clustering framework. Furthermore, we collaborated with maintainers of popular tools for metagenomic community profiling^49,50^ to offer built-in support for UHGV, facilitating its integration into existing workflows.

## Methods

### Integration and quality control of virome databases

We obtained virus genomes and genome fragments derived from 12 recently published phage genome catalogues (Table S1), including the MGV^13^, GPD^14^, GVD^12^, GFD^15^, CHVD^16^, COPSAC^23^, HuVirDB^11^, IMG/VR v4^22^, mMGE^19^, DEVoC^17^, as well as viruses previously identified in metagenomic assemblies of faecal samples from 5,742 diverse human populations^18^ and from the Hadza hunter-gatherers^24^. For the MGV dataset, we included the 189,680 contigs with >50% completeness described in the original publication and an additional 304,160 viral contigs longer than 10 kb with ≤50% completeness that were identified in the original study but not included in the public data release. For IMG/VR, we ignored contigs already present in MGV.

The original datasets were integrated using a consistent pipeline for virus genome verification and quality assessment, which combined multiple tools: geNomad^51^ (version 1.1.0), viralVerify^52^ (version 1.1), and CheckV^53^ (version 1.0.1, database version 1.4). First, all putative viral genomes were processed using geNomad, and sequences with an aggregate virus score lower than 0.7 were discarded, except those matching *Inoviridae* marker genes (see below). Proviruses flanked by host genes were excised using both geNomad and CheckV. Sequences passing geNomad detection were then assigned to confidence tiers using a signature-based scoring system. We defined three viral signatures: (1) geNomad aggregated virus score > 0.95, (2) classification as “Virus” by viralVerify with score > 15, or (3) presence of at least 3 geNomad virus hallmark genes, or at least 3 CheckV viral genes with a 3× excess of viral over host genes. Four non-viral signatures were also defined: (1) classification as “Plasmid” or “Chromosome” by viralVerify, (2) at least 2 CheckV host genes with host genes outnumbering viral genes, (3) at least one geNomad plasmid hallmark gene, or (4) at least one conjugation gene. A net viral score was calculated for each contig by subtracting the number of non-viral signatures (maximum 4) from the number of viral signatures (maximum 3). Finally, sequences were assigned to one of three confidence tiers: non-viral (score ≤ −1), uncertain virus (score of 0 or 1), or confident virus (score ≥ 2).

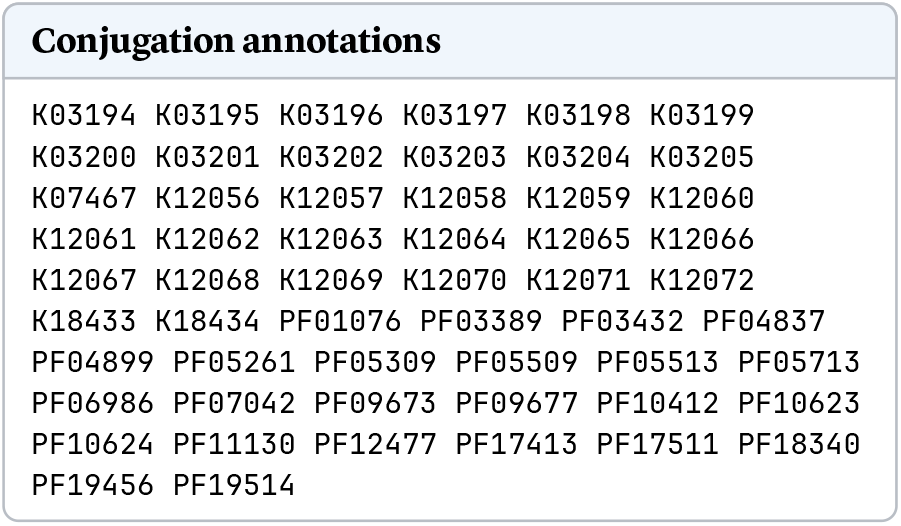

An exception was made for sequences assigned to a known ICTV family, in which case we only required a net viral score ≥ 1 to be labelled as a confident virus. We also made an exception for putative *Inoviridae*, which were often mislabelled as plasmids or left unlabelled by the viral classifiers. To confidently identify *Inoviridae*, we looked for contigs that contained (1) the Zot protein (Pfam accession: PF05707), (2) at least 5 genes that matched taxonomically-informative markers for *Inoviridae* from the geNomad database, and (3) a genome length between 4,500 and 12,500 bp. The Zot protein is homologous to morphogenesis proteins that are widely distributed in *Inoviridae* and has been used as a marker for this taxon^54.^ Any contig labelled as non-viral was removed, while uncertain and confident viruses were retained in the database.

To classify sequences into quality tiers, the presence of direct terminal repeats (DTRs) and CheckV completeness estimates were evaluated^53.^ Genomes with DTRs were considered complete if they had >80% CheckV-estimated completeness and, for *Caudoviricetes*, a genome size >17 kb^55.^ DTRs were trimmed from the right side of viral genomes. For genomes without DTRs, completeness was estimated with CheckV, using a reference database that was supplemented with the DTR-containing genomes. To further polish the dataset, we removed: (1) contigs shorter than 10 kb with ≤50% estimated genome completeness, (2) contigs with >500 ambiguous bases (“Ns”) or where Ns comprised >1% of the contig, (3) contigs with an average 21-mer frequency >1.2, which can result from a large duplicated region or highly repetitive elements^53^, (4) contigs that were >1.5× longer than the expected genome size based on CheckV, which can indicate the presence of tandem integrated proviruses or a provirus flanked by a large host region was not removed by CheckV.

### Clustering genomes into species-level viral OTUs

Average nucleotide identity (ANI) and alignment fraction (AF) were estimated for genome pairs from an all-versus-all BLAST search^56^ (version 2.10.1+, parameters: `-task megablast -outfmt ‘6 std qlen slen’ -max_target_seqs 10000`), processed using a custom script (available at https://bitbucket.org/berkeleylab/checkv/src/master/scripts/anicalc.py). Analysis of 1,000 randomly selected high-quality and complete genomes showed that our ANI and AF estimates were highly concordant with those from MUMMer4^57^ (Pearson’s *ρ* for ANI and AF: 0.980 and 0.948, respectively) and were robust to circularly permuted genomes. Following MIUViG^26^ guidelines, we retained genome connections with ≥95% ANI over ≥85% of the shorter sequence, and calculated inter-genome similarities as (ANI × AF)/100.

We used an iterative strategy to cluster the 874,103 UHGV genomes into 168,569 species-level vOTUs. First, we clustered the complete genomes with DTRs into *de novo* vOTUs. Next, linear sequences with >90% completeness were assigned to existing vOTUs with the remaining unassigned contigs clustered *de novo*. Lastly, linear contigs with ≤90% completeness were assigned to an existing vOTUs with the remaining unassigned contigs clustered *de novo*. All clustering was performed with MCL^58^ (version 14-137) using default parameters. Our novel, iterative clustering strategy was designed to prevent short fragments and proviruses integrated in tandem from bridging together complete genomes of distantly related viruses.

Last, we selected a single representative genome per species-level vOTU. The longest complete genome was selected, so long as it was in the upper 50th percentile of genome size for the vOTU, otherwise, we chose the linear genome that was closest to the expected genome size based on CheckV and contained the most CheckV viral genes. Our strategy for picking a representative genome was designed to select the most complete genome without biasing selection towards short genome fragments with DTRs or very long sequences that are flanked by bacterial DNA.

### Clustering genomes into taxonomy-like higher ranking clusters

Representative genomes from species-level vOTUs were further grouped into broader clusters in a hierarchical system similar to traditional taxonomy, with ranks based on varying degrees of a genome-wide proteomic similarity measurement^23^. This score ranges from 0 to 1 and was computed as follows: (1) perform an all-versus-all alignment of protein sequences from viral genomes using DIAMOND^59^ (version 2.0.9.147, parameters: `-very-sensitive -k 100000 -e 1e-3`), (2) calculate the cumulative bit score across best blast hits between each pair of viral genomes, (3) calculate a normalisation constant for each genome, obtained from the cumulative bit score of best hits from a self alignment, and (4) divide the bit score from (2) by (3) to obtain the final proteomic similarity score.

Clustering began by removing edges between genome pairs with low similarity, grouping genomes into family-level clusters, and pruning the graph to retain only intra-family connections. This was repeated for subfamily, genus, and subgenus ranks using progressively higher similarity thresholds. At each rank, edges below the threshold were removed, genomes were clustered, and inter-cluster connections were pruned. Clustering was performed using the MCL algorithm with default parameters at all levels. The minimum proteomic similarity at each level was chosen in order to maximise agreement with ICTV taxa at that rank^60^ and previously proposed *Crassvirales* groups^38^, quantified using the adjusted mutual information metric. The following cutoffs were selected: 80% (vSUBGEN), 65% (vGENUS), 32% (vSUBFAM), and 5.5% (vFAM). To limit false positives, we only formed higher ranking clusters using the high-quality viral genomes. Taxonomic coherence was maintained by ensuring that all genomes classified within the same taxon at any given rank also shared taxa at all higher ranks.

### Taxonomic assignment

Taxonomic assignment of vOTUs to viral taxa was conducted using genome-wide proteomic similarity to reference genomes and taxonomically informative protein profiles from geNomad. A reference database was created by combining phage genomes from the INPHARED database (retrieved in Apr-2022)^61^, with taxonomic lineages modified to include only taxa recognised in ICTV’s VMR 19/MSL 37, and metagenomic phage genomes from newly proposed taxa prevalent in the human gut. These included the five *Crassvirales* groups from Yutin *et al*.^38^ and the *Flandersviridae, Gratiaviridae*, and *Quim byviridae* families from Benler *et al*.^18^ The proteomic similarity of vOTU representatives to reference genomes was calculated as described for genome clustering, requiring a minimum similarity of 5.5%. Phages were then assigned to the lineage of their closest reference match, with taxa transferred down to a rank determined by the similarity level (cutoffs: 65% for genus, 32% for subfamily, and 5.5% for family). vOTUs lacking sufficient similarity to any reference genome were assigned to taxa inferred by geNomad using protein alignments to taxonomically informative MMseqs2 profiles.

### Genome annotation

Protein coding genes were predicted using prodigal-gv^51,62^ (version 2.10.0, parameters: `-p meta`), which identified more than 98% of alternative coded phages identified in our previous study^13^. To limit mispredictions of alternative genetic codes, we only report genomes that are at least 10 kb as recoded. tRNAs were predicted using tRNAscan-SE^63^ (version 2.0.9), and DGR systems were identified with DGRscan^64^.

Functional annotations were obtained using multiple of tools and databases: eggNOG-mapper^65^ (version 2.1.6), InterProScan^66^ (version 5.66-98.0, parameters: `-appl HAMAP`), PHROG^67^ (release 3), Pfam^68^ (release 36.0), NCBIfam^69^ (release 13.0), KOfam^70^ (version 2022-02-01), UniRef90^71^ (release Feb 2022), PADLOC-DB^72^ (release 1.3.0), dbAPIS^73^ (2023-09-19 update), and AcrCatalog^74^. COG (release COG2020)^75^ and HAMAP^76^ identifiers were obtained from the outputs of eggNOG-mapper and InterProScan, respectively. PHROG, Pfam, NCBIfam, KOfam, PADLOC-DB, and dbAPIS homology searches were performed using PyHMMER^77,78^ (version 0.5.0, hmmsearch parameters: `bit_cutoffs=“gathering”` for Pfam and NCBIfam, `E=1e-5` for the remaining databases). Searches against UniRef90 and AcrCatalog were conducted with DIAMOND (parameters: `-query-cover 50 -subject-cover 50`). For Pfam, multiple matches to a single protein were allowed if the matches did not overlap or if the overlap covered at most 20% of the shortest domain’s length. For other databases, only the best hit was accepted.

Virus lifestyle was predicted using BACPHLIP^79^ (version 0.9.6), which uses conserved protein domains to predict lifestyle via a random forest classifier. A virus was classified as temperate if the BACPHLIP score was higher than 0.5, if it contained an integrase from the PHROG database, or if it was excised from a larger contig by geNomad, otherwise the virus was classified as virulent. As BACPHLIP was trained on complete genomes, it may not be accurate for genome fragments that are missing integrases and other genes.

### Protein clustering

Using MMseqs2^80^ (version 14.7e284), protein sequences were clustered in two steps. First, an all-vs-all alignment with a minimum 80% bi-directional coverage was performed, followed by greedy set cover clustering (parameters: `cluster -s 7.0 -c 0.8 -cluster-steps 3 -cluster-reassign 1 -kmer-per-seq 50`). Next, multiple sequence alignments were generated for each cluster using MMseqs2′s centre star alignment, converted to protein profiles, and used to obtain consensus sequences. These consensus sequences were then employed in a profile-vs-protein search with a minimum 90% bi-directional coverage (parameters: `search -s 8.0 -c 0.9`), which guided the second clustering step.

### Structure prediction, clustering, and annotation

Multiple sequence alignments were generated using FAMSA^81^ (version 2.2.2-7eb7612) for protein clusters containing at least 14 unique sequences. To improve structure prediction performance, these alignments were enriched to increase sequence diversity using HHblits^82^ (version 3.3.0, parameters: `-n 3`), with UniRef30^83^ (release 2023-02) and the UHGV protein clusters alignments used as target databases.

Structure prediction was performed with ColabFold^84^ via the LocalColabFold installer (version 1.5.2, parameters: `-max-seq 512 -max-extra-seq 1024`, available at https://github.com/YoshitakaMo/localcolabfold) for clusters with representative sequences up to 2,000 residues. Poor quality predictions (pLDDT < 50) were discarded and the remaining structures were functionally annotated using Foldseek^85^ (commit dffdf78) and clustered. For clustering, Reseek^86^ (version 2.5, parameters: `-sensitive`) was used for all-versus-all structure alignment, generating a weighted graph where edges, weighted by alignment quality, connect structure pairs with bidirectional coverage ≥ 90% and alignment quality ≥ 0.85. This graph was then clustered with pyLeiden^87^ (version 0.1.2, parameters: `-r 0.44`, available at https://github.com/apcamargo/pyleiden). Protein domain segmentation was achieved using Merizo^88^ (commit fbb38be, parameters: `-plddt_filter 50 -conf_filter 0.75`).

To functionally annotate UHGV structures, we utilised Foldseek to search reference structure databases from Pfam, NCBIfam, HAMAP (release 13-Sep-2023), and COG (2020 release). These reference structures were predicted from multiple sequence alignments provided by the respective sources, employing the same routine applied to UHGV protein clusters. Structural alignment cutoffs were set as follows: target coverage ≥ 0.4, probability ≥ 0.9, and E-value ≤ 1.0. These thresholds were determined by comparing Foldseek results to gold standard annotations using the Pfam and NCBIfam reference databases. The gold standard was derived from hmmsearch-based annotations of 1,500 randomly selected protein clusters, each containing at least 250 members with at least 80% sharing the same annotation. Optimal cutoffs were determined through a grid search maximising precision^2^ × recall, where precision was defined as the fraction of structural hits where Foldseek annotations matched hmmsearch annotations, and recall as the fraction of hmmsearch annotations correctly identified by Foldseek. The same set of thresholds was used to assess similarity between UHGV structures and PDB structures (retrieved on 06-Dec-2023).

### Building a comprehensive prokaryote genome database

A novel human gut prokaryote genome database was constructed to enable viral host assignment and metagenomic read recruitment analysis. This database comprises 464,666 prokaryotic genomes, including 286,387 from the UHGG v2^89^, 54,779 from a study of the gut microbiome of Hadza hunter-gatherer and rural Nepali individuals^24^, and 123,500 representative genomes from GTDB r207 that were either found in the HumGut database^90^ or detected in at least one of 1,000 healthy human gut metagenomes using the `screen` command of Mash^91^ (version 2.3).

The genomes were clustered and assigned to specieslevel clusters in three stages. First, we grouped the 464,666 genomes into 45,725 clusters at 97.5% ANI using dRep^92^ (version 3.3.0, parameters: `-sa 0.975`). Second, representative genomes for each cluster were selected as the ones with the highest genome score, computed as: completeness − 5 × contamination + 0.5 × log_10_ (N50) +log_10_(genome length) + 2 × (centrality ™ 0.95) × 100). Here centrality is defined as the mean Mash distance between a given genome and all other genomes within the same species-level group. Taxonomy was then assigned to each representative genome using GTDB-Tk^93^ (version 2.1.0, database release r207), resulting in 40,385 assignments to known species from the GTDB which were then extended to the remaining cluster members. Third, representative genomes from the 97.5% ANI clusters that were not assigned to a species by GTDB-Tk were clustered at 95% ANI using dRep (parameters: `-sa 0.95`) into *de novo* operational species. At the end of this process, 6,754 species-level clusters were obtained, with 449,161 genomes assigned to known species from the GTDB and 15,505 from *de novo* clustering.

### Host assignment and host range breadth computation

To assign vOTU representative genomes from the UHGV to prokaryotic host taxa, we used our newly constructed prokaryote genome database. We employed two methods to establish connections between viruses and hosts: aligning CRISPR spacers extracted from the prokaryotic genomes to viral sequences, and using k-mer matching to directly identify viruses within the genomes of their hosts.

CRT^94^ (version 1.1) and PILER-CR^95^ (version 1.06) were used to identify a total of 5,318,089 CRISPR spacers from the prokaryotic genomes. To minimise false positives^96^, we removed CRISPR arrays with: (1) fewer than 3 spacers, (2) an average repeat length < 22 bp or >55 bp, (3) an average spacer length < 25 bp, or (4) containing non-conserved repeats (<85% average identity to the consensus repeat), and (5) individual spacers < 25 bp or containing ambiguous bases (e.g. NNN). CRISPR spacers were matched to the vOTU representatives using BLAST (version 2.10.1+, parameters: `-task megablast -max_target_seqs 10000 -word_size=8 -dust no`) and we retained spacer alignments with a maximum of one mismatch or gap that covered at least 95% of the spacer’s length. For host assignment through detection in prokaryotic genomes, we used PHIST^97^ (version 1.0.0, parameters: `-k 30`) for alignment-free k-mer matching. To accurately capture viral host range and avoid spurious connections, we considered all virushost connections but required ≥20% of the k-mers of a virus genome to be detected in the genome of its host.

Host taxon assignment was then performed independently for the CRISPR and PHIST methods. We assigned vOTU representatives to the host taxon at the lowest taxonomic rank that comprised at least 70% of the connections using the `find_majority_vote` of the taxopy library (version 0.11.0, parameters: `fraction=0.7`). To integrate methods and obtain the final host prediction, we used the host prediction method resulting in the greater number of host connections for each vOTU representative.

Host range breadth was quantified by measuring distances between host genomes within the bacterial phylogeny using SuchTree^98^ (version 1.0). We retrieved the GTDB r207 Bacteria phylogeny, rooted it to place *Patescibacteria* as a sister group to all other bacteria, and performed taxonomic-rank normalisation^99^ using PhyloRank’s `outlier` command (version 0.1.12) to ensure cophenetic distances between species were between 0 and 1. We then computed vOTU host range breadth as the median cophenetic distance across pairs of host species to which the vOTU had at least one connection, weighting each pair’s distance by the product of the number of phage-host links for the two species.

### Identification of functions associated with increased host range breadth

To identify functions associated with increased HRB, we compared the HRB of vOTUs with each protein annotation to those without it. Only high-quality and complete genomes were used, and annotations were required to be detected in at least 970 vOTU representatives (approximately 2.5% of the vOTUs). To eliminate highly collinear annotations, we calculated the variance inflation factor (VIF) for each and removed those with a VIF greater than 5. This process was performed iteratively, removing the least frequent annotation with a high VIF until no annotation exceeded the threshold. To account for confounding factors, an ordinary least squares regression was conducted with host range breadth as the dependent variable, while the number of host genomes, predicted lifestyle, and common host taxa (*Actinomycetia, Bacilli, Bacteroidia, Clostridia, Gammaproteobacteria*, and *Negativicutes*) were used as independent variables. Residuals from this model were used to identify annotations significantly associated with host range breadth by comparing genomes with and without each annotation using two-sided t-tests with unequal variances. The maximum false discovery rate, computed using the Benjamini-Yekutieli procedure, was set to 5%. Further statistical assessment of the associations was conducted using phylogenetic regression models (`phylolm` function, parameters: `model = “kappa”`), as implemented in the phylolm^100 l^ibrary (version 2.6.2). In addition to the phylogeny-based estimate, host range breadth was quantified using an alternative metric that uses Shannon entropy, which measures the distribution of host genera and families linked to each vOTU by the number of connections to different taxa at the genus and family levels.

### *Caudoviricetes* phylogenetic reconstruction

A phylogenetic tree of *Caudoviricetes* phages was reconstructed using concatenated multiple sequence alignments (MSAs) of the ATPase and endonuclease domains of the large terminase subunit (TerL), the major capsid protein, and the portal protein. To enhance orthologue detection sensitivity, we developed the diversify-pfam tool (version 0.1.0, available at https://github.com/apcamargo/diversify-pfam), which generates protein hidden Markov models (HMMs) to detect target domains. diversify-pfam enriched MSAs from 13 selected Pfam, NCBIfam, and HAMAP accessions (see list below) using HHblits with the UniRef30 and BFD databases^101^. It then clustered the enriched MSAs into sub-MSAs of closely related sequences, which were then used to generate HMMER-formatted HMMs.

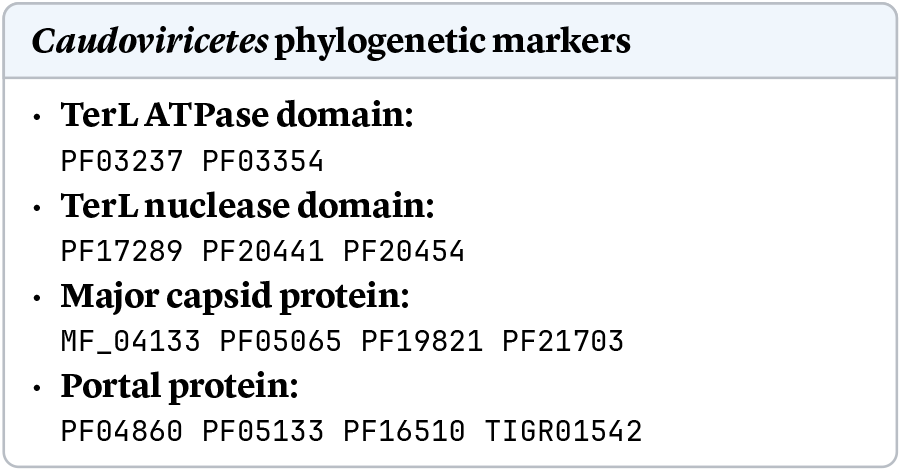

To detect target proteins within UHGV genomes, we employed PyHMMER’s hmmsearch function (version 0.5.0, parameters: `T=10` for *TerL* domains, `T=25` for other genes) to align proteins to HMMs, requiring at least 60% HMM coverage. Aligned protein regions were extracted and regions with the same annotation were aligned with Muscle^102^ (version 5.1, parameters: `-super5`). Proteins matching HMMs from different annotations were assigned to the HMM with the highest bit score. The resulting 13 MSAs were trimmed with ClipKIT^103^ (version 2.3.0), concatenated, and filtered to remove concatenated alignments with gaps across more than 90% of their extension. Finally, FastTree^104^ (version 2.1.11, parameters: `-wag -gamma`) was used to reconstruct the phylogeny.

### Profiling of viruses and prokaryotes in worldwide human gut metagenomes

We used a read recruitment approach to assess the prevalence and abundance of viruses and prokaryotes in human gut microbiomes worldwide. We selected 1,798 bulk metagenomes (each with at least 5 million reads) representing diverse human populations. Additionally, we processed 673 viromes, including all publicly available human gut viromes on NCBI (as of 01-Feb-2022) with at least 500,000 reads and a viral enrichment score >10, as determined by ViromeQC^105^ (version 1.0.1). Reads were mapped to medium-quality or better vOTU representatives using Bowtie 2^106^ (version 2.4.5). Unmapped reads were then mapped to our collection of prokaryote genomes using the same method. Virus and prokaryote BAM files were profiled using CoverM^107^ (version 0.6.1, parameters: `-m mean trimmed_mean covered_bases length count - min-covered-fraction 0`). Viral contigs were considered detected if at least 75% of their bases were covered, while prokaryote genomes were considered detected if at least 50% of their bases were covered. Metagenomes were classified into enterotypes using Enterotyper^108^ (commit befca5c), with genus-level aggregated relative abundances of prokaryotic genomes as input to the PAM clustering algorithm with the 2-enterotype model.

Hyperprevalent vOTUs were defined as those detected in ≥50 samples across three or more countries. To assess their distribution across lifestyles (“industrialised”, “hunter-gatherers”, “transitional”), we used a re-sampling approach to address sampling biases, selecting up to 40 samples per lifestyle (no more than 10 per country) and averaging detections across 30 iterations. vOTUs were assigned to a lifestyle if more than two thirds of the detections occurred in that lifestyle. If over three-fourths of detections were shared between two lifestyles, vOTUs were categorised accordingly (e.g., “hunter-gatherers and transitional”). vOTUs not meeting these criteria were classified as “all lifestyles”. Lifestyle classifications were assigned based on manual review of sample metadata and associated literature.

To compute vOTU enterotype enrichment (the log ratio of the average relative abundances within each enterotype) and identify hyperprevalent vOTUs, we used a subset of the data. Samples were excluded if they (1) were subjected to amplification (“MDA”, “WGA”, “WTA”, “phi29”, or “amplification” mentioned in the library preparation metadata, or ≥50% of the detected vOTUs assigned to the *Monodnaviria* realm), (2) had shallow sequencing depth (“pyrosequencing” or “genome analyzer” mentioned in the sequencing method metadata, or <15 distinct vOTUs detected), or (3) targeted RNA (>1% of the detected vOTUs were assigned to the *Riboviria* realm).

### Detection of single-nucleotide and single-codon variants

Intra-population virus genome diversity was assessed by detection of single-nucleotide variants and single-codon variants^109^ from read mapping data using anvi’o^110^ (version 8). BAM files produced with Bowtie 2 (see section above) were processed using the `anvi-profile` (parameters: `-min-coverage-for-variability 10 -profile-SCVs`) and `anvi-gen-variability-profile` (parameters: `-engine NT`) commands to identify variant sites. To filter sequencing artifacts, rare variants were removed if the ratio of base variant frequency to the most common base at that position did not exceed a dynamic cutoff: 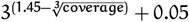 retention of rare alleles at high sequencing depths. For single-codon variant detection, the output of anvi-gen-variabilityprofile (parameters: `-engine CDN`) was processed to remove spurious variants using the same dynamic cutoff. To focus on within-population polymorphisms, we retained only sites and codons exhibiting multiple alleles detected within a single sample. The strength of selection acting on polymorphic codons was quantified by comparing the potentials for nonsynonymous and synonymous substitutions (see Supplementary Note 3 for details).

## Data availability

Genome and protein sequences, functional annotations, predicted protein structures, genome metadata, read mapping data, and single-nucleotide variant data are available for download at https://uhgv.jgi.doe.gov/downloads (Zenodo DOI: 10.5281/zenodo.17402089).

## Code availability

The toolkit that enables the assignment of external viral genomes to UHGV clusters within the UHGV taxonomy-like genome clustering framework is available as open-source software at https://github.com/snayfach/UHGV-classifier (Zenodo DOI: 10.5281/zenodo.17418882). It includes both the software and database required for classifying viral genomes and evaluating their novelty relative to UHGV reference genomes.

## Acknowledgements

The work conducted by the U.S. Department of Energy Joint Genome Institute (https://ror.org/04xm1d337), a DOE Office of Science User Facility, is supported by the Office of Science of the U.S. Department of Energy operated under Contract No. DE-AC02-05CH11231. This research used resources provided by the National Energy Research Scientific Computing Center (NERSC), a Department of Energy User Facility (project m306-2024), and by the São Paulo Research Foundation (FAPESP, grant 2021/10577-0). A.P.C., S.R., and N.C.K. were supported in part by the National Institutes of Health (NIH, grant 5U01DE034196-02). F.A.B. and G.A.P. were supported by the Hellenic Foundation for Research and Innovation (H.F.R.I.) under the “Third Call for H.F.R.I. Research Projects to support faculty members and researchers” [23592 - EMIS-SION].

## References

1. Lynch, S. V. & Pedersen, O. The Human Intestinal Microbiome in Health and Disease. New England Journal of Medicine 375, 2369–2379 (2016).

2. Kurilovich, E. & Geva-Zatorsky, N. Effects of bacteriophages on gut microbiome functionality. Gut Microbes 17, 2481178 (2025).

3. Nepal, R., Houtak, G., Wormald, P.-J., Psaltis, A. J. & Vreugde, S. Prophage: a crucial catalyst in infectious disease modulation. The Lancet Microbe 3, e162–e163 (2022).

4. Norman, J. M. et al. Disease-Specific Alterations in the Enteric Virome in Inflammatory Bowel Disease. Cell 160, 447–460 (2015).

5. De Jonge, P. A. et al. Gut virome profiling identifies a widespread bacteriophage family associated with metabolic syndrome. Nature Communications 13, 3594 (2022).

6. Yang, K. et al. Alterations in the Gut Virome in Obesity and Type 2 Diabetes Mellitus. Gastroenterology 161, 1257–1269.e13 (2021).

7. Gordillo Altamirano, F. L. & Barr, J. J. Phage Therapy in the Postantibiotic Era. Clinical Microbiology Reviews 32, e66–18 (2019).

8. Gontijo, M. T. P., Jorge, G. P. & Brocchi, M. Current Status of Endolysin-Based Treatments against Gram-Negative Bacteria. Antibiotics 10, 1143 (2021).

9. Dutilh, B. E. et al. A highly abundant bacteriophage discovered in the unknown sequences of human faecal metagenomes. Nature Communications 5, 4498 (2014).

10. Devoto, A. E. et al. Megaphages infect Prevotella and variants are widespread in gut microbiomes. Nature Microbiology 4, 693–700 (2019).

11. Soto-Perez, P. et al. CRISPR-Cas System of a Prevalent Human Gut Bacterium Reveals Hyper-targeting against Phages in a Human Virome Catalog. Cell Host & Microbe 26, 325–335.e5 (2019).

12. Gregory, A. C. et al. The Gut Virome Database Reveals Age-Dependent Patterns of Virome Diversity in the Human Gut. Cell Host & Microbe 28, 724–740.e8 (2020).

13. Nayfach, S. et al. Metagenomic compendium of 189,680 DNA viruses from the human gut microbiome. Nature Microbiology 6, 960–970 (2021).

14. Camarillo-Guerrero, L. F., Almeida, A., Rangel-Pineros, G., Finn, R. D. & Lawley, T. D. Massive expansion of human gut bacteriophage diversity. Cell 184, 1098– 1109.e9 (2021).

15. Garmaeva, S. et al. Stability of the human gut virome and effect of gluten-free diet. Cell Reports 35, 109132 (2021).

16. Tisza, M. J. & Buck, C. B. A catalog of tens of thousands of viruses from human metagenomes reveals hidden associations with chronic diseases. Proceedings of the National Academy of Sciences 118, e2023202118 (2021).

17. Van Espen, L. et al. A Previously Undescribed Highly Prevalent Phage Identified in a Danish Enteric Virome Catalog. mSystems 6, e382–21 (2021).

18. Benler, S. et al. Thousands of previously unknown phages discovered in whole-community human gut metagenomes. Microbiome 9, 78 (2021).

19. Lai, S. et al. mMGE: a database for human metagenomic extrachromosomal mobile genetic elements. Nucleic Acids Research 49, D783–D791 (2021).

20. Tomofuji, Y. et al. Prokaryotic and viral genomes recovered from 787 Japanese gut metagenomes revealed microbial features linked to diets, populations, and diseases. Cell Genomics 2, 100219 (2022).

21. Nishijima, S. et al. Extensive gut virome variation and its associations with host and environmental factors in a population-level cohort. Nature Communications 13, 5252 (2022).

22. Camargo, A. P. et al. IMG/VR v4: an expanded database of uncultivated virus genomes within a framework of extensive functional, taxonomic, and ecological metadata. Nucleic Acids Research 51, D733–D743 (2023).

23. Shah, S. A. et al. Expanding known viral diversity in the healthy infant gut. Nature Microbiology 8, 986–998 (2023).

24. Carter, M. M. et al. Ultra-deep sequencing of Hadza hunter-gatherers recovers vanishing gut microbes. Cell 186, 3111–3124.e13 (2023).

25. Dutilh, B. E. et al. Perspective on taxonomic classification of uncultivated viruses. Current Opinion in Virology 51, 207–215 (2021).

26. Roux, S. et al. Minimum Information about an Uncultivated Virus Genome (MIUViG). Nature Biotechnology 37, 29–37 (2019).

27. Chao, A. et al. Rarefaction and extrapolation of phylogenetic diversity. Methods in Ecology and Evolution 6, 380–388 (2015).

28. International Committee on Taxonomy of Viruses Executive Committee et al. The new scope of virus taxonomy: partitioning the virosphere into 15 hierarchical ranks. Nature Microbiology 5, 668–674 (2020).

29. Simmonds, P. et al. Virus taxonomy in the age of metagenomics. Nature Reviews Microbiology 15, 161–168 (2017).

30. Bin Jang, H. et al. Taxonomic assignment of uncultivated prokaryotic virus genomes is enabled by gene-sharing networks. Nature Biotechnology 37, 632–639 (2019).

31. Tamburini, F. B. et al. Short- and long-read metagenomics of urban and rural South African gut microbiomes reveal a transitional composition and undescribed taxa. Nature Communications 13, 926 (2022).

32. Lou, Y. C. et al. Infant gut DNA bacteriophage strain persistence during the first 3 years of life. Cell Host & Microbe 32, 35–47.e6 (2024).

33. Grigson, S. R., Giles, S. K., Edwards, R. A. & Papudeshi, B. Knowing and Naming: Phage Annotation and Nomenclature for Phage Therapy. Clinical Infectious Diseases 77, S352–S359 (2023).

34. Varadi, M. et al. AlphaFold Protein Structure Database: massively expanding the structural coverage of protein-sequence space with high-accuracy models. Nucleic Acids Research 50, D439–D444 (2022).

35. Illergård, K., Ardell, D. H. & Elofsson, A. Structure is three to ten times more conserved than sequence—A study of structural response in protein cores. Proteins: Structure, Function, and Bioinformatics 77, 499–508 (2009).

36. Smug, B. J., Szczepaniak, K., Rocha, E. P. C., Dunin-Horkawicz, S. & Mostowy, R. J. Ongoing shuffling of protein fragments diversifies core viral functions linked to interactions with bacterial hosts. Nature Communications 14, 7460 (2023).

37. Arumugam, M. et al. Enterotypes of the human gut microbiome. Nature 473, 174–180 (2011).

38. Yutin, N. et al. Analysis of metagenome-assembled viral genomes from the human gut reveals diverse putative CrAss-like phages with unique genomic features. Nature Communications 12, 1044 (2021).

39. Klimasauskas, S., Kumar, S., Roberts, R. J. & Cheng, X. Hhal methyltransferase flips its target base out of the DNA helix. Cell 76, 357–369 (1994).

40. Liu, M. et al. Reverse transcriptase-mediated tropism switching in \textit{Bordetella} bacteriophage. Science 295, 2091–2094 (2002).

41. Wu, L. et al. Diversity-generating retroelements: natural variation, classification and evolution inferred from a large-scale genomic survey. Nucleic Acids Research 46, 11–24 (2018).

42. Kiefl, E. et al. Structure-informed microbial population genetics elucidate selective pressures that shape protein evolution. Science Advances 9, eabq4632 (2023).

43. Murphy, J., Mahony, J., Ainsworth, S., Nauta, A. & Van Sinderen, D. Bacteriophage Orphan DNA Methyltransferases: Insights from Their Bacterial Origin, Function, and Occurrence. Applied and Environmental Microbiology 79, 7547–7555 (2013).

44. Shaw, L. P., Rocha, E. P. C. & MacLean, R. C. Restriction-modification systems have shaped the evolution and distribution of plasmids across bacteria. Nucleic Acids Research 51, 6806–6818 (2023).

45. Li, S. & Waters, R. \textit{Escherichia coli} strains lacking protein HU are UV sensitive due to a role for HU in homologous recombination. Journal of Bacteriology 180, 3750–3756 (1998).

46. Rigden, D. J., Jedrzejas, M. J. & Galperin, M. Y. Amidase domains from bacterial and phage autolysins define a family of γ-d,l-glutamate-specific amidohydrolases. Trends in Biochemical Sciences 28, 230–234 (2003).

47. Griffin, M. E., Klupt, S., Espinosa, J. & Hang, H. C. Peptidoglycan NlpC/P60 peptidases in bacterial physiology and host interactions. Cell Chemical Biology 30, 436– 456 (2023).

48. Oechslin, F., Zhu, X., Dion, M. B., Shi, R. & Moineau, S. Phage endolysins are adapted to specific hosts and are evolutionarily dynamic. PLOS Biology 20, e3001740 (2022).

49. Pinto, Y., Chakraborty, M., Jain, N. & Bhatt, A. S. Phage-inclusive profiling of human gut microbiomes with Phanta. Nature Biotechnology 42, 651–662 (2024).

50. Shaw, J. & Yu, Y. W. Rapid species-level metagenome profiling and containment estimation with sylph. Nature Biotechnology 43, 1348–1359 (2025).

51. Camargo, A. P. et al. Identification of mobile genetic elements with geNomad. Nature Biotechnology 42, 1303– 1312 (2024).

52. Antipov, D., Raiko, M., Lapidus, A. & Pevzner, P. A. MetaviralSPAdes: assembly of viruses from metagenomic data. Bioinformatics 36, 4126–4129 (2020).

53. Nayfach, S. et al. CheckV assesses the quality and completeness of metagenome-assembled viral genomes. Nature Biotechnology 39, 578–585 (2021).

54. Roux, S. et al. Cryptic inoviruses revealed as pervasive in bacteria and archaea across Earth’s biomes. Nature Microbiology 4, 1895–1906 (2019).

55. Luque, A., Benler, S., Lee, D. Y., Brown, C. & White, S. The Missing Tailed Phages: Prediction of Small Capsid Candidates. Microorganisms 8, 1944 (2020).

56. Chen, Y., Ye, W., Zhang, Y. & Xu, Y. High speed BLASTN: an accelerated MegaBLAST search tool. Nucleic Acids Research 43, 7762–7768 (2015).

57. Marçais, G. et al. MUMmer4: A fast and versatile genome alignment system. PLOS Computational Biology 14, e1005944 (2018).

58. Van Dongen, S. Graph Clustering Via a Discrete Uncoupling Process. SIAM Journal on Matrix Analysis and Applications 30, 121–141 (2008).

59. Buchfink, B., Reuter, K. & Drost, H.-G. Sensitive protein alignments at tree-of-life scale using DIAMOND. Nature Methods 18, 366–368 (2021).

60. Lefkowitz, E. J. et al. Virus taxonomy: the database of the International Committee on Taxonomy of Viruses (ICTV). Nucleic Acids Research 46, D708–D717 (2018).

61. Cook, R. et al. INfrastructure for a PHAge REference Database: Identification of Large-Scale Biases in the Current Collection of Cultured Phage Genomes. PHAGE 2, 214–223 (2021).

62. Hyatt, D. et al. Prodigal: prokaryotic gene recognition and translation initiation site identification. BMC Bioinformatics 11, 119 (2010).

63. Chan, P. P., Lin, B. Y., Mak, A. J. & Lowe, T. M. tRNAscanSE 2.0: improved detection and functional classification of transfer RNA genes. Nucleic Acids Research 49, 9077– 9096 (2021).

64. Ye, Y. Identification of Diversity-Generating Retroelements in Human Microbiomes. International Journal of Molecular Sciences 15, 14234–14246 (2014).

65. Cantalapiedra, C. P., Hernández-Plaza, A., Letunic, I., Bork, P. & Huerta-Cepas, J. eggNOG-mapper v2: Functional Annotation, Orthology Assignments, and Domain Prediction at the Metagenomic Scale. Molecular Biology and Evolution 38, 5825–5829 (2021).

66. Jones, P. et al. InterProScan 5: genome-scale protein function classification. Bioinformatics 30, 1236–1240 (2014).

67. Terzian, P. et al. PHROG: families of prokaryotic virus proteins clustered using remote homology. NAR Genomics and Bioinformatics 3, lqab67 (2021).

68. Paysan-Lafosse, T. et al. The Pfam protein families database: embracing AI/ML. Nucleic Acids Research 53, D523–D534 (2025).

69. Li, W. et al. RefSeq: expanding the Prokaryotic Genome Annotation Pipeline reach with protein family model curation. Nucleic Acids Research 49, D1020–D1028 (2021).

70. Aramaki, T. et al. KofamKOALA: KEGG Ortholog assignment based on profile HMM and adaptive score threshold. Bioinformatics 36, 2251–2252 (2020).

71. Suzek, B. E. et al. UniRef clusters: a comprehensive and scalable alternative for improving sequence similarity searches. Bioinformatics 31, 926–932 (2015).

72. Payne, L. J. et al. Identification and classification of antiviral defence systems in bacteria and archaea with PADLOC reveals new system types. Nucleic Acids Re search 49, 10868–10878 (2021).

73. Yan, Y., Zheng, J., Zhang, X. & Yin, Y. dbAPIS: a database of a ntip rokaryotic i mmune s ystem genes. Nucleic Acids Research 52, D419–D425 (2024).

74. Gussow, A. B. et al. Machine-learning approach expands the repertoire of anti-CRISPR protein families. Nature Communications 11, 3784 (2020).

75. Galperin, M. Y. et al. COG database update 2024. Nucleic Acids Research 53, D356–D363 (2025).

76. Pedruzzi, I. et al. HAMAP in 2015: updates to the protein family classification and annotation system. Nucleic Acids Research 43, D1064–D1070 (2015).

77. Larralde, M. & Zeller, G. PyHMMER: a Python library binding to HMMER for efficient sequence analysis. Bioinformatics 39, btad214 (2023).

78. Sean R. Eddy, Travis Wheeler & Nick Carter. HMMER: biosequence analysis using profile hidden Markov models. http://hmmer.org/.

79. Hockenberry, A. J. & Wilke, C. O. BACPHLIP: predicting bacteriophage lifestyle from conserved protein domains. PeerJ 9, e11396 (2021).

80. Steinegger, M. & Söding, J. MMseqs2 enables sensitive protein sequence searching for the analysis of massive data sets. Nature Biotechnology 35, 1026–1028 (2017).

81. Deorowicz, S., Debudaj-Grabysz, A. & Gudyś, A. FAMSA: Fast and accurate multiple sequence alignment of huge protein families. Scientific Reports 6, 33964 (2016).

82. Steinegger, M. et al. HH-suite3 for fast remote homology detection and deep protein annotation. BMC Bioinformatics 20, 473 (2019).

83. Mirdita, M. et al. Uniclust databases of clustered and deeply annotated protein sequences and alignments. Nucleic Acids Research 45, D170–D176 (2017).

84. Mirdita, M. et al. ColabFold: making protein folding accessible to all. Nature Methods 19, 679–682 (2022).

85. Van Kempen, M. et al. Fast and accurate protein structure search with Foldseek. Nature Biotechnology 42, 243– 246 (2024).

86. Edgar, R. C. Protein structure alignment by Reseek improves sensitivity to remote homologs. Bioinformatics 40, btae687 (2024).

87. Traag, V. A., Waltman, L. & Van Eck, N. J. From Louvain to Leiden: guaranteeing well-connected communities. Scientific Reports 9, 5233 (2019).

88. Lau, A. M., Kandathil, S. M. & Jones, D. T. Merizo: a rapid and accurate protein domain segmentation method using invariant point attention. Nature Communications 14, 8445 (2023).

89. Almeida, A. et al. A unified catalog of 204,938 reference genomes from the human gut microbiome. Nature Biotechnology 39, 105–114 (2021).

90. Hiseni, P., Rudi, K., Wilson, R. C., Hegge, F. T. & Snipen, L. HumGut: a comprehensive human gut prokaryotic genomes collection filtered by metagenome data. Microbiome 9, 165 (2021).

91. Ondov, B. D. et al. Mash Screen: high-throughput sequence containment estimation for genome discovery. Genome Biology 20, 232 (2019).

92. Olm, M. R., Brown, C. T., Brooks, B. & Banfield, J. F. dRep: a tool for fast and accurate genomic comparisons that enables improved genome recovery from metagenomes through de-replication. The ISME Journal 11, 2864–2868 (2017).

93. Chaumeil, P.-A., Mussig, A. J., Hugenholtz, P. & Parks, D. H. GTDB-Tk v2: memory friendly classification with the genome taxonomy database. Bioinformatics 38, 5315– 5316 (2022).

94. Bland, C. et al. CRISPR Recognition Tool (CRT): a tool for automatic detection of clustered regularly interspaced palindromic repeats. BMC Bioinformatics 8, 209 (2007).

95. Edgar, R. C. PILER-CR: Fast and accurate identification of CRISPR repeats. BMC Bioinformatics 8, 18 (2007).

96. Mitrofanov, A. et al. CRISPRidentify: identification of CRISPR arrays using machine learning approach. Nucleic Acids Research 49, e20–e20 (2021).

97. Zielezinski, A., Deorowicz, S. & Gudyś, A. PHIST: fast and accurate prediction of prokaryotic hosts from metagenomic viral sequences. Bioinformatics 38, 1447– 1449 (2022).

98. Y. Neches, R. & Scott, C. SuchTree: Fast, thread-safe computations with phylogenetic trees. Journal of Open Source Software 3, 678 (2018).

99. Parks, D. H. et al. A standardized bacterial taxonomy based on genome phylogeny substantially revises the tree of life. Nature Biotechnology 36, 996–1004 (2018).

100. Tung Ho, L. S. & Ané, C. A Linear-Time Algorithm for Gaussian and Non-Gaussian Trait Evolution Models. Systematic Biology 63, 397–408 (2014).

101. Steinegger, M. & Söding, J. Clustering huge protein sequence sets in linear time. Nature Communications 9, 2542 (2018).

102. Edgar, R. C. Muscle5: High-accuracy alignment ensembles enable unbiased assessments of sequence homology and phylogeny. Nature Communications 13, 6968 (2022).

103. Steenwyk, J. L., Buida, T. J., Li, Y.Shen, X.-X. & Rokas, A. ClipKIT: A multiple sequence alignment trimming software for accurate phylogenomic inference. PLOS Biology 18, e3001007 (2020).

104. Price, M. N., Dehal, P. S. & Arkin, A. P. FastTree 2 – Approximately Maximum-Likelihood Trees for Large Alignments. PLOS One 5, e9490 (2010).

105. Zolfo, M. et al. Detecting contamination in viromes using ViromeQC. Nature Biotechnology 37, 1408–1412 (2019).

106. Langmead, B. & Salzberg, S. L. Fast gapped-read alignment with Bowtie 2. Nature Methods 9, 357–359 (2012).

107. Aroney, S. T. N. et al. CoverM: read alignment statistics for metagenomics. Bioinformatics 41, btaf147 (2025).

108. Keller, M. I. et al. Refined Enterotyping Reveals Dysbiosis in Global Fecal Metagenomes. bioRxiv (2024) doi:10.1101/2024.08.13.607711.

109. Delmont, T. O. et al. Single-amino acid variants reveal evolutionary processes that shape the biogeography of a global SAR11 subclade. eLife 8, e46497 (2019).

110. Eren, A. M. et al. Community-led, integrated, reproducible multi-omics with anvi’o. Nature Microbiology 6, 3–6 (2020).

